# COLOR VISION UNDER BLUR: IMPLICATIONS FOR PERCEPTION AND EVOLUTION

**DOI:** 10.64898/2026.03.31.715493

**Authors:** Nil Altinordu, Geoffrey Boynton, Ione Fine

## Abstract

Color is a prominent feature of visual experience, yet humans can recognize objects easily and accurately from grayscale images. We examined whether color becomes more useful when spatial information is degraded due to blurring. Participants viewed naturalistic scenes in color or grayscale, and reported whether a named target object was present across a range of blur levels that simulated optical defocus from 0–8 diopters. With unblurred images, performance did not differ between color and grayscale conditions, but as blur increased, recognition accuracy declined. Color provided a modest but reliable advantage at higher levels of blur, suggesting that color becomes increasingly useful when optical quality is degraded. We hypothesize that the evolutionary shift towards trichromacy may have been partially driven by the need to compensate for optical degradation due to aging and/or accumulated light exposure.

## INTRODUCTION

Color is a prominent feature of visual experience that is supported by specialized mechanisms throughout the visual system. Physiological and psychophysical studies have long demonstrated distinct chromatic pathways in early vision (Gegenfurtner & Kiper, 2003; Livingstone & Hubel, 1987). Consistent with this, neuroimaging studies show that color contributes to object perception through neural networks that are partly distinct from those processing shape and form (Bannert & Bartels, 2013; Brouwer & Heeger, 2009; Lafer-Sousa et al., 2016).

Despite this, the functional role of color for object recognition remains debated. Objects can be recognized easily and accurately even when color information is absent. People have little difficulty following black-and-white films, and individuals with congenital color vision deficiencies typically function in everyday environments without obvious recognition impairments (Steward & Cole, 1989). Recognition of objects in line drawings or grayscale photographs, approaches the accuracy of recognition with full-color images (Biederman & Ju, 1988; Rossion & Pourtois, 2004). These observations raise a basic question: what is color actually good for in object recognition?

Color does seem to facilitate recognition when it is strongly associated with an object’s identity. Objects with high color diagnosticity (e.g., bananas or pumpkins) are recognized more quickly when presented in their canonical colors than when presented in grayscale or incongruent colors (Tanaka & Presnell, 1999). Similar effects have been observed across a variety of tasks, including naming and verification (Therriault et al., 2009), and canonical color can sometimes facilitate recognition when spatial information is limited (Lewis et al., 2013). However, relatively few objects are highly color diagnostic, and the behavioral advantages reported in these studies are typically modest.

The present study examined the role of color in scene-based object detection under defocus blur. Participants viewed photographic scenes and reported whether a target object was present. Images were presented either in color or grayscale and were blurred to simulate increasing levels of optical defocus.

Understanding how blur and color interact in naturalistic vision has become increasingly important with the advent of sight-restoration technologies. These devices currently only restore coarse spatial vision with no useful chromatic information. Moreover, clinical measures of vision (e.g., Snellen or perimetry) do not assess the contributions of color information. When evaluating the likely effectiveness of sight recovery devices, it is important to understand how consequential the absence of color information may be under conditions of degraded spatial information.

Our use of naturalistic scenes differs from most previous studies, which have tended to focus on the recognition of objects presented in isolation against uniform backgrounds. In natural vision, objects appear within complex scenes and must be detected amidst visual clutter. Scene context, segmentation, and figure–ground organization therefore become important components of the task. Because color can provide useful cues for segmentation and grouping, its contribution may be particularly relevant in scene-based object detection, where the visual system must separate objects from their backgrounds before recognition can occur.

Similarly, the interaction between color and blur has received surprisingly little attention. Wurm et al. (1993) examined recognition of fruit images under optical blur and found main effects of color and blur but no interaction between them. However, these studies relied on stimuli that were either highly color diagnostic or presented against simplified backgrounds, leaving open the question of how color and blur interact in natural scenes.

It seems plausible that color becomes more useful in complex scenes under conditions of blur. Defocus attenuates high spatial frequencies, weakening the contour and shape cues that normally support rapid object recognition. Natural images contain both luminance-defined and chromatically defined edges, and these signals are not entirely redundant. Chromatic edges can exist where luminance edges are weak or absent, providing information about object boundaries that is partly independent of luminance contrast (Fine et al., 2003; Hansen & Gegenfurtner, 2009). Moreover, color information is shifted towards lower spatial frequencies (Parraga et al., 1998) and sensitivity to these chromatic signals is low pass (Mullen, 1985); thus color may be informative under blurred conditions where fine spatial detail is unavailable.

Our primary finding is that that color provides little advantage when images are well focused, and becomes increasingly useful under conditions of heavy blur. This finding provides a potential novel explanation for the development of trichromacy. Prevailing accounts of primate trichromacy emphasize its role in detecting fruit or young leaves against foliage (Mollon, 1989; Osorio & Vorobyev, 1996; Sumner & Mollon, 2000). Longer-lived monkeys in the wild, especially those in high light environments might well have degraded optical quality later in life, due to presbyopia and/or a loss of dichromatic sensitivity along the blue-yellow dimension due to yellowing of the lens or cataracts. If chromatic signals are particularly useful under degraded optical conditions, then trichromacy, especially in the form of enhanced discrimination between longer wavelengths, may help compensate for age and light-exposure related optical degradation. We find that ecological proxies for lifetime light exposure successfully predict the presence of routine trichromacy across diurnal primate species.

## METHODS

We carried out three experiments using the same stimuli and object-detection task. Experiment 1 measured baseline performance with grayscale and color unblurred images with unlimited viewing time, Experiment 2 tested the effects of color and blur with unlimited viewing time, and Experiment 3 tested the same conditions as Experiment 2 under time-limited viewing.

Across all three experiments, 57 participants (38 female, 19 male) with normal or corrected-to-normal visual acuity were recruited through word of mouth at the University of Washington. Participants provided informed consent in accordance with procedures approved by the University of Washington Human Subjects Division and were compensated for their time. All participants were fluent in English, and individuals with known color vision deficiencies were excluded. Participants were distributed across experiments as follows: 12 in Experiment 1, 27 in Experiment 2, and 18 in Experiment 3. Each experiment used a unique set of participants.

Stimuli were presented on a Cambridge Research Systems Display++ LCD monitor (1920 × 1080 resolution, 120 Hz refresh rate) with a physical screen size of 70 × 39 cm. The monitor was gamma-linearized prior to testing. The viewing distance was approximately 107 cm with free viewing, resulting in stimuli subtending approximately 20° × 32° of visual angle. The monitor was the only source of illumination in the testing room. Experimental code was written in MATLAB using Psychtoolbox (Brainard, 1997; Pelli, 1997).

### Stimuli

Stimuli consisted of 177 (Experiment 1) images of naturalistic scenes, of which 140 were used in Experiments 2 and 3. Images were sourced from two sources: the Berlin Object in Scene (BOiS) database (https://info.ni.tu-berlin.de/photodb/) and Adobe Stock Images.

The BOiS dataset (74 images) consisted of real-world photographs containing paired images in which a target object is either present or absent in a plausible scene location. In our task, participants reported whether the target object was present in the scene, and either the object-present or object-absent image was presented on each trial. Adobe Stock images (70 images) consisted of real-world photographs and realistic AI-generated scenes. For these images, either an existing object in the scene was designated as the target object, or a plausible target object was selected that could realistically appear in the scene but was not present.

Images were cropped to 4041 × 2590 pixels and presented on the central portion of the display, subtending approximately 20° × 32° of visual angle. Grayscale versions of each image were generated in MATLAB. Blurred stimuli were created using a disk filter to simulate defocus blur ranging from 0 to 8 diopters in 1-diopter increments (Strasburger et al., 2018). Example stimuli are shown in Figure 1.

**Figure 1.**
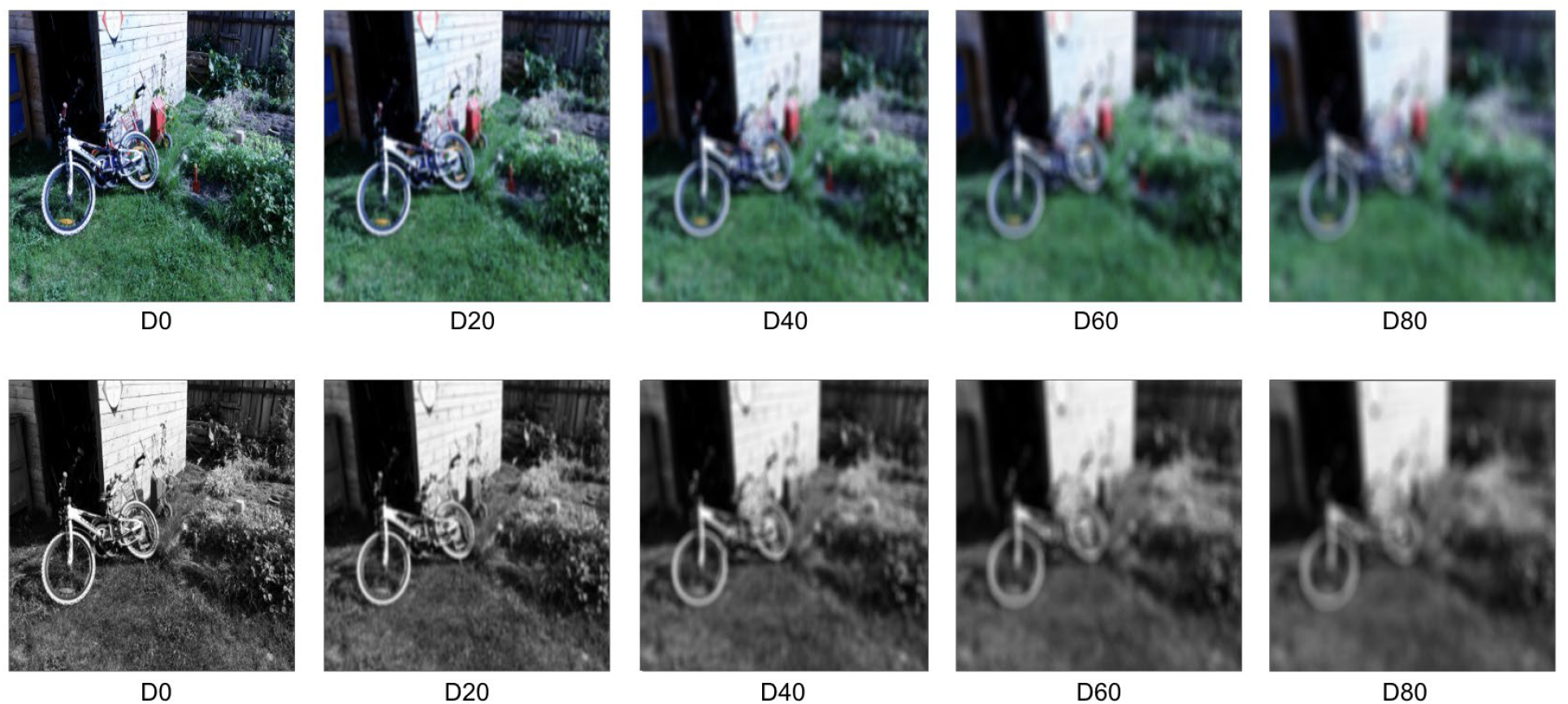
Example color and grayscale stimuli at 0, 2, 4, 6, and 8 diopters of simulated blur. Object names were verified with two native American English speakers to ensure that each target object name was an unambiguous and commonly used label. All experimental participants were fluent English speakers.

### Experiment 1: Baseline performance with unblurred images

Experiment 1 measured baseline object detection performance for unblurred images. The experiment used a 2 (Color: color, grayscale) × 2 (Object presence: present, absent) within-subject design. Twelve participants (5 female) completed this experiment. Each participant completed 36 trials per condition (144 trials total). Image assignment and trial order were pseudorandomized across participants.

Each trial began with the name of the target object presented in white text on a gray background for 1 s. This was immediately followed by the scene image. Participants responded with a key press to indicate whether the target object was present or absent. The scene remained visible until a response was made. Participants received auditory feedback (high or low beep) indicating whether their response was correct. Both accuracy and reaction times (RTs) were recorded.

Although objects were pre-screened to ensure that their names were easily understood, for some scenes/objects participants consistently showed poor accuracy. In Experiments 2 and 3 we excluded any images for which fewer than 75% of participants responded correctly in Experiment 1. This ensured that all the stimuli used in these two experiments were consistently recognizable under unblurred conditions.

### Experiment 2: Effects of blur with unlimited viewing

Experiment 2 examined whether color improves object detection in natural scenes when images are degraded by blur. Twenty-seven participants (18 female) completed this experiment. The experiment used a 2 (Color: color, grayscale) × 2 (Object presence: present, absent) × 9 (Blur level) within-subject design. Each participant completed 7 trials per condition (144 trials total).

The procedure was identical to Experiment 1 except that the images were presented with varying levels of simulated blur. Both accuracy and reaction times were recorded.

### Experiment 3: Effects of blur under time-limited viewing

Experiment 3 tested whether the effects observed in Experiment 2 would also occur under time-limited viewing conditions. Eighteen participants (15 female) completed this experiment.

The design and stimuli were identical to those used in Experiment 2. The only difference was that scene presentation was time limited. Instead of remaining visible until a response was made, the scene was presented for 2.5 seconds (65% of responses in Experiment 1 were made within 2.5s), after which it was replaced by a mask.

### Statistical Methods – Psychophysics

Trial-level accuracy (correct vs. incorrect) was analyzed with R using a generalized linear mixed-effects model with a binomial error distribution and logit link function implemented in lme4. We tested for the main effects of color and blur and their interaction, with subject as a random effect factor. Model parameters were estimated using maximum likelihood, and statistical significance of fixed effects was evaluated using Type II Wald χ^2^ tests (car::Anova).

For Experiments 2 and 3, we conducted post-hoc comparisons to examine the effect of color separately across three levels of blur: low (0-2 diopters), medium (3-5 diopters) and high (6-8 diopters). Estimated marginal means were computed from the fitted model using R’s emmeans package, and pairwise contrasts comparing color versus grayscale were evaluated within each level. P-values were adjusted across the three levels using the Holm–Bonferroni procedure to control the familywise error rate.

Standard logistic regression models were fit using glm with a binomial error distribution.

### Modeling: The Relationship between Trichromacy and Lifetime Light Exposure

All statistical analyses were conducted at the species level using a dataset of diurnal primates containing information on color vision, diet, lifespan, geographic distribution, and habitat use (see Appendix 1). Data processing and analysis were implemented in R. Analyses were restricted to species with complete data for all variables included in a given model (complete-case filtering).

Routine trichromacy ( *Trichromacy*_*i*_ )was treated as a binary response variable. Percent frugivory (the consumption of fruit as a primary food source, *Frugivory*_*i*_)was derived from dietary data and standardised to a continuous percentage variable. Maximum Lifespan (*Lifespan*_*i*_ )was obtained by combining available longevity estimates within the dataset.

A proxy for ambient short wave/ultra-violet light exposure as a function of latitude was approximated as *UV*_*i*_= cos(|∅|), where ∅_*i*_ is absolute latitude in degrees, for each species, *i*.

Habitat use was represented using a categorical canopy variable with three levels:

0 = non-arboreal (terrestrial)

1 = high canopy

2 = low canopy or mixed canopy use

Since broadleaf tree canopies reduce UV exposure by somewhere between 70-95% (Parker et al., 2019; Qi et al., 2010; Sivarajah et al., 2020) a canopy multiplier was applied to represent reduced light exposure in arboreal environments. This was done by creating a Canopy exposure multiplier for each species (*Canopy*_*i*_)that represented ecological light exposure as follows (results were robust to the exact choice of multiplier:

1.0 for non-arboreal species

0.5 for high-canopy species

0.1 for low-canopy or understory species

Lifetime light exposure was then calculated as:

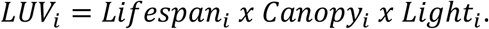

Continuous predictors were standardised (z-transformed) prior to analysis to allow comparison of effect sizes. We fit a standard binomial logistic regression model treating species as independent observations:

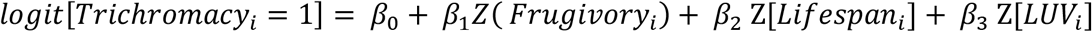

Reduced models (Frugivory only, Lifespan only, Lifetime Light Exposure only, and Canopy only) were fitted to evaluate the independent contribution of each predictor. Model fit was compared using Akaike Information Criterion (AIC). Coefficients were interpreted as log-odds and converted to odds ratios for reporting.

### Software

All analyses were conducted in R (version 4.5.2). Databases and models are provided at https://github.com/VisCog/color_blur

## RESULTS

### Experiment 1: Baseline performance with unblurred images

Figure 2 Panel A shows accuracy across grayscale and color images for Experiment 1. Mean accuracy was 85.6% for color images and 85.9% for grayscale images. Logistic mixed-effects models revealed no significant effect of color on accuracy (χ^2^(1) = 0.035, p = .851). Figure 2B shows RTs, which also did not differ across color and grayscale images. Mean RTs were 2.01s for grayscale and 1.99s for color. Logistic mixed-effects models revealed no significant effect of color on RTs (χ^2^(1) = 0.368, p = .544). These results indicate that color does not improve object recognition speed or accuracy when scenes are sharply focused.

**Figure 2.**
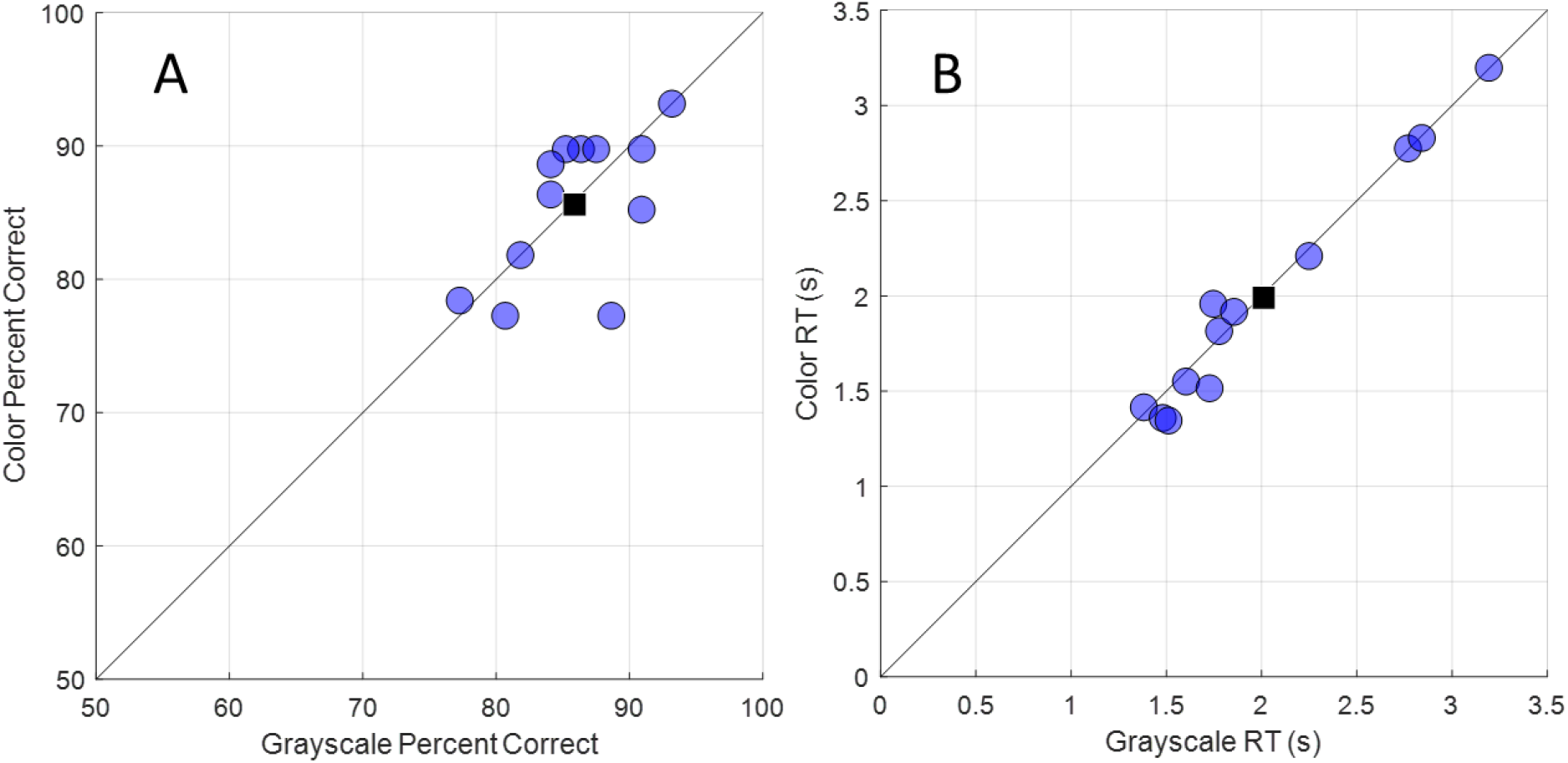
**Panel A.** Percent correct for object recognition with no blurring for individual subjects in Experiment 1. Filled blue symbols show grayscale performance on the x-axis and color performance on the y-axis for each subject. Mean performance across subjects is shown as the black square. **Panel B**. RTs for object recognition without blurring from individual subjects in Experiment 1. Filled blue symbols show grayscale performance on the x-axis and color performance on the y-axis for each subject. Mean RTs across subjects are shown as the black square.

Although objects were pre-screened to ensure that their names were easily understood, recognition performance on some images was relatively poor. To ensure reliable recognition in the blur experiments, we excluded any images for which fewer than 75% of participants responded correctly across both the color and gray conditions of Experiment 1. This eliminated 27 of the 177 images (∼15%). As a result, 140 unique images were used in Experiments 2 and 3.

### Experiment 2: Effects of blur with unlimited viewing

Experiment 2 examined whether color improves object detection when scenes are degraded by blur. Results, averaged across participants, are shown in Figure 3A.

**Figure 3.**
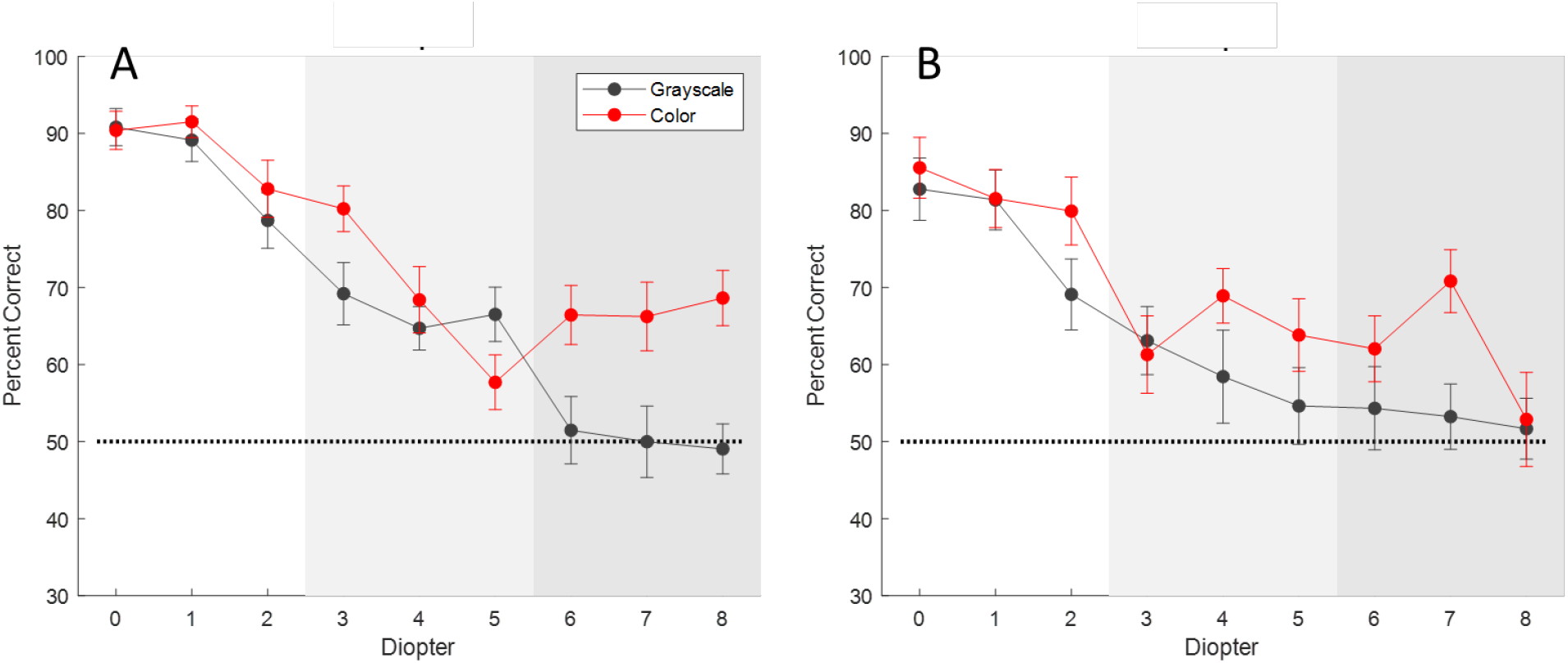
Results from Experiment 2 (Panel A) and Experiment 3 (Panel B). Object recognition percent correct, averaged across subjects, for grayscale and color images across 9 levels of blur. Error bars were computed as within-subject standard errors using the Cousineau–Morey method applied to subject-level proportion-correct values. This normalization removes between-subject differences in overall performance, such that the error bars reflect variability across conditions within subjects.

Statistical inference was based on a binomial generalized linear mixed-effects model fit to trial-level responses. A Type II Wald χ^2^ analysis revealed a significant main effect of Blur, χ^2^(8) = 207.13, p < .001 - accuracy declined as blur increased. There was also a significant main effect of Color, χ^2^(1) = 15.49, p < .001, with higher overall accuracy for color than grayscale images.

The Blur × Color interaction was significant, χ^2^(8) = 19.64, p = .012, indicating that the effect of color depended on the level of blur. Inspection of the data showed that performance for color and grayscale images was similar at low blur levels but diverged as blur increased.

We conducted post-hoc comparisons to examine the effect of color at low (0-2 diopters), medium (3-5 diopters) and high (6-8 diopters) levels of blur. After Holm-Bonferroni correction for multiple comparisons across the three blur levels, color did not have a significant advantage at the lower levels of blur (0-2 diopters: z = .631, p = .528, 3-5 diopters: z = .756, p = .450), but did improve object recognition at the highest levels of blur (6-9 diopters: z = 4.898, p <.001).

Analysis of RT data found that RTs were *fastest* for the highest blur levels, especially for the grayscale condition. This is probably because performance in these conditions were near chance, and subjects may have ‘given up’ and responded quickly. Because this made the RT data uninterpretable, RT results are not shown for Experiments 2 or 3.

### Experiment 3: Effects of blur under time-limited viewing

Experiment 3 examined whether the color advantage observed in Experiment 2 would also occur when viewing time was limited to 2.5 seconds (the 65% percentile reaction time of Experiment 1), followed by a mask. Results are shown in Figure 3B.

As in Experiment 2, there was a significant main effect of blur, χ^2^(8) = 88.06, p < .001, indicating that accuracy decreased as blur increased. There was also a significant main effect of color, χ^2^(1) = 7.74, p = .005, with higher accuracy for color images than grayscale images. In contrast to Experiment 2, there was not a significant Color x Blur interaction χ^2^(8) = 7.1967, p = 0.516. However, this lack of an interaction seemed to be due to performance being at chance for both color and blur at the highest blur level (8 diopters).

We conducted the same post-hoc comparisons as in Experiment 2, examining the effect of color at low (0-2 diopters), medium (3-5 diopters) and high (6-8 diopters) levels of blur. This analysis showed very similar results. After Holm-Bonferroni correction for multiple comparisons across the three blur levels, color did not have a significant advantage at the lowest levels of blur (0-2 diopters: z = 1.186, p = .235, 3-5 diopters: z = 1.338, p = .181), but color did improve object recognition at the highest levels of blur (6-9 diopters: z = 2.338, p <.019).

### Results: The Relationship between Trichromacy and Lifetime Light exposure

A total of 143 diurnal primate species were included in the non-phylogenetic analysis, of which 84 (58.7%) were classified as routine trichromats.

For the full binomial logistic regression model, both lifespan and lifetime light exposure were significant positive predictors of routine trichromacy. Specifically, routine trichromacy was positively associated with log-transformed lifespan (β = 0.716 ± 0.278 SE, *p* = 0.010) and with lifetime light exposure (β = 0.717 ± 0.267 SE, *p* = 0.007). In contrast, the frugivory predictor was non-significant; percent frugivory was not significantly associated with trichromacy (β = 0.067 ±0.206 SE, *p* = 0.743). The full model substantially improved fit relative to the null model (residual deviance = 163.64 vs. 193.85) and had the lowest AIC among all models (AIC = 171.64).

When considered individually, lifetime light exposure was a strong predictor of routine trichromacy (β = 0.964 ± 0.268 SE, *p* < 0.001), producing a large reduction in deviance (172.08) and a low AIC (176.08). Similarly, lifespan alone was strongly associated with trichromacy (β = 1.052 ± 0.259 SE, *p* < 0.001; AIC = 177.53).

Canopy use was also a significant predictor (β = −1.460 ± 0.388 SE, *p* < 0.001), indicating that species occupying lower canopy strata were less likely to exhibit routine trichromacy. However, the canopy-only model provided a poorer fit than models including lifetime light exposure or lifespan (AIC = 177.24).

Frugivory showed no significant effect (β = 0.242 ± 0.176 SE, *p* = 0.169) and did not improve model fit relative to the null (AIC = 195.91).

Thus, our statistical model shows that animals with longer lifetimes, and high lifetime light exposure have a higher probability of trichromacy. However, it is important to note that this analysis does not account for the considerable impact of shared ancestry, as discussed below.

## Discussion

The goal of the present study was to determine whether color contributes to object recognition when spatial information is degraded by blur. Across experiments, we found that color provided no benefit under sharply focused conditions (Experiment 1) but conferred a modest (∼10-15%) yet reliable advantage at the highest levels of blur (Experiments 2 and 3). This pattern supports the hypothesis that chromatic information becomes increasingly useful as luminance-defined shape cues are degraded.

Our hypothesis that color may compensate for blur, is supported by the observation that chromatic and luminance signals differ systematically in their spatial frequency content. Natural image analyses show that color carries information at coarser spatial scales(Parraga et al., 1998) and human chromatic contrast sensitivity is biased toward lower spatial frequencies (Mullen, 1985). Because defocus blur selectively attenuates high spatial frequencies, it disproportionately disrupts luminance-based shape information while leaving chromatic signals relatively intact. Our observed benefit of colour under blur may not simply be an additive effect, but might reflect a shift in the relative reliability of visual cues. As high-frequency luminance information is lost, recognition may depend more heavily on chromatic signals that remain informative at lower spatial scales.

Our use of naturalistic scenes allowed us to examine the role of color in a relatively naturalistic object recognition task which is reasonably reflective of how objects are recognized in the real world. However, our use of a complex task does mean that color may have contributed to task performance in multiple ways.

In cluttered environments, object recognition depends not only on identifying features but also on separating objects from their backgrounds in the context of search. In the absence of blur, luminance-defined contours and texture provide sufficient information for identifying objects, leaving little opportunity for color to improve performance (Biederman & Ju, 1988). As blur increases, high-frequency spatial information is progressively lost, weakening the shape cues that normally support recognition, especially within the context of cluttered natural scenes. Under these conditions, chromatic information, which is partly independent of luminance boundaries (Fine et al., 2003; Hansen & Gegenfurtner, 2009), may provide complementary information to support object segmentation and recognition.

Blur may also increase the reliance on stored priors about object appearance. Color is an important component of stored knowledge about objects, and many objects have characteristic colors that can facilitate retrieval of object representations (Bramao et al., 2011; Tanaka & Presnell, 1999). Under degraded viewing conditions, our observers may have relied more heavily on learned associations to infer object identity from partial visual information.

These findings have important implications for visual prostheses and other sight-restoration technologies. Current prosthetic vision systems provide limited spatial resolution and lack meaningful chromatic information. Our results suggest that color does contribute to object recognition when spatial content is degraded – making it important to factor in the absence of chromatic information when assessing the utility of these devices.

Finally, our statistical modelling suggests that one environmental driver of the development of trichromacy may be the need to compensate for optical deterioration. The dominant hypothesis is that trichromacy evolved to facilitate detection of fruit or young leaves against foliage (Mollon, 1989; Osorio & Vorobyev, 1996). Our results suggest a non-exclusive alternative possibility –that chromatic vision provides a compensatory advantage under conditions of optical degradation arising from ageing or cumulative light exposure. Aging in high light conditions is likely to cause optical degradation that includes increased blur due to presbyopia and lens yellowing/cataracts that will disproportionately attenuate short-wavelength transmission. Under such conditions, the ability to make chromatic discrimination within the longer wavelength spectrum might plausibly provide an evolutionary advantage. It should be noted that this should be considered a complementary rather than a competitive hypothesis – the spectral sensitivity conferred by trichromacy might be optimized for foraging with poor optics.

Our analysis does not account for the considerable impact of shared ancestry. Because trichromacy has evolved only a small number of times in primates, phylogenetic constraints limit the ability to draw strong causal inferences. In Old World primates, trichromacy arose through a duplication of an X-linked opsin gene, creating separate medium- and long-wavelength pigments (Carvalho et al., 2017; Hunt et al., 1998). In contrast, most New World primates retain a single polymorphic gene that, for heterozygous females, produces sex-linked trichromacy (Jacobs et al., 1996; Mollon, 1989). The key exception is the howler monkey, which independently evolved Old World–style trichromacy through a separate duplication event. This convergent evolution has typically interpreted as having been driven by the need to detect reddish or yellowish food against green foliage or fresh green leaves from mature leaves (Mollon, 1989; Osorio & Vorobyev, 1996) However, this pattern of evolutionary events is equally consistent with the notion that trichromacy evolved to compensate for high lifetime light exposure. Old World monkeys have long lifespans and tend to live within high light environments. Howler monkeys, even though they are arboreal, have relatively long lifespans and, critically, have a strong preference for the upper middle to high canopy.

Overall, our findings demonstrate that color provides a modest but reliable benefit to object recognition when spatial information is degraded, supporting the idea that chromatic cues become increasingly important as luminance-defined shape information is lost. Extending these findings to an evolutionary context, our analyses suggest that optical deterioration due to old age and/or lifetime light exposure may represent a plausible selective pressure for the emergence of trichromacy in long-lived species, especially those exposed to high light environments.

