## Appendix 1 for "COLOR VISION UNDER BLUR: IMPLICATIONS FOR PERCEPTION AND EVOLUTION"

We developed a curated primate trait spreadsheet containing species-level information on colour vision, diet, longevity, activity pattern, geographic range, and foraging stratum. We began with a database that included lifespan, trichromacy and other variables derived from PanTHERIA, EltonTraits, and AnAge, together with rule-based classifications of routine trichromacy and activity pattern. Species names were standardized and duplicate entries removed. Analyses were restricted to taxa included in the non-nocturnal build of the spreadsheet.

Routine trichromacy was coded as a binary variable, where species with routine trichromatic vision were assigned 1 and all other taxa were assigned 0. Percent frugivory was taken from the EltonTraits-derived fruit diet variable. Geographic light environment was represented using absolute midpoint latitude.

We created an ecological canopy variable with three categories: 0 = non-arboreal, 1 = high canopy, and 2 = low canopy or mixed canopy use. This variable was assigned using a simple ecological rule set: selected genera with well-known terrestrial or upper-canopy habits were manually mapped to canopy categories, and all remaining taxa were classified using arboreality information already present in the dataset.

Species names were standardised before phylogenetic matching, and complete-case filtering was applied at analysis time rather than during initial dataset construction.
